# OrthoHMM: Improved Inference of Ortholog Groups using Hidden Markov Models

**DOI:** 10.1101/2024.12.07.627370

**Authors:** Jacob L Steenwyk, Thomas J. Buida, Antonis Rokas, Nicole King

## Abstract

Accurate orthology inference is essential for comparative genomics and phylogenomics. However, orthology inference is challenged by sequence divergence, which is pronounced among anciently diverged organisms. We present OrthoHMM, an algorithm that infers orthologous gene groups using Hidden Markov Models parameterized from substitution matrices, which enables better detection of remote homologs. Benchmarking indicates OrthoHMM outperforms currently available methods; for example, using a curated set of Bilaterian orthogroups, OrthoHMM showed a 10.3 – 138.9% improvement in precision. Rank-based benchmarking using Bilaterian orthogroups and a novel dataset of orthogroups from organisms in three major eukaryotic kingdoms revealed OrthoHMM had the best overall performance (6.7 – 97.8% overall improvement). These findings suggest that Hidden Markov Models improve orthogroup inference.

## Main text

Global orthology inference aims to group genes from different organisms into sets of orthologs. This analysis is often an essential step for comparative genomics and Tree of Life inference^1,2^. However, global orthology inference is notoriously challenging due to, for example, variation in sequence divergence, duplication followed by divergence, and gene loss^3^.

Current methods frequently use networks—wherein nodes represent genes and edges reflect sequence similarity—to group genes into clusters of putative orthologs, termed orthogroups^4–7^. To do so, all-by-all sequence comparisons are made using proteomes from a set of taxa, and recent advances have focused on how the resulting sequence similarity scores are used to infer network edges^6^. Top-performing algorithms determine edge weights from database-independent similarity measures (e.g., bitscores calculated by BLAST) and apply phylogenetic corrections to account for increased sequence divergence across greater evolutionary distances^4,5^. These algorithms typically use BLAST, DIAMOND, or MMseqs for quantifying sequence similarity^8–10^. Profile Hidden Markov Models (HMMs), which use probabilistic modelling and are known to be more sensitive, can be helpful in cases where sequence divergence is pronounced, such as among anciently diverged organisms and rapidly evolving gene families^11^. Other methods infer orthologs using alternative information, such as phylogeny-aware methods^12^, but these require additional computation and may not be suitable for large-scale ortholog inference.

We introduce OrthoHMM, an algorithm that infers global orthology using phylogenetically corrected sequence similarity scores calculated using HMMs. OrthoHMM uses a special implementation of HMMs wherein the profile is parameterized using an amino acid substitution matrix rather than a multiple sequence alignment of representative protein sequences. OrthoHMM takes as input a directory of FASTA files as well as optional arguments that allow fine-tuned user control. OrthoHMM then internally conducts the all-by-all comparisons, network construction, and clustering (Fig. 1; algorithm details are available in the supplementary text). To facilitate downstream analyses, OrthoHMM outputs summary files—such as a taxon-by-orthogroup matrix of gene counts and lists of genes per orthogroup—and directories of all orthogroups and single-copy orthogroups in FASTA format.

**Figure 1.**
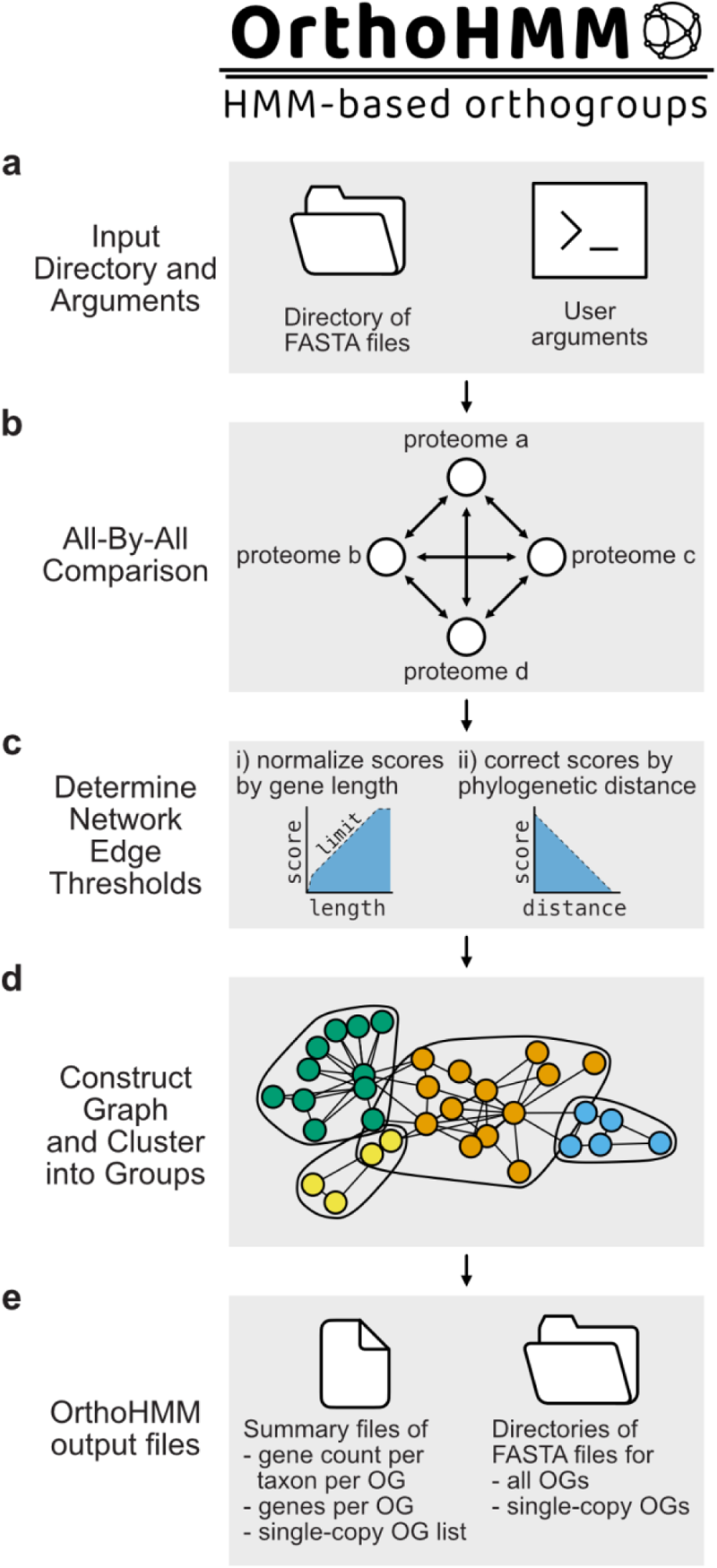
An overview of the OrthoHMM workflow. (a) OrthoHMM takes the path to a directory of FASTA protein sequence files and optional arguments, such as an e-value threshold or amino acid substitution matrix, as input for building profile HMMs. (b-e) Thereafter, OrthoHMM internally handles all subsequent steps. (b) Specifically, OrthoHMM conducts an all-by-all sequence comparison and (c) corrects the resulting sequence similarity scores for gene length and phylogenetic distance biases. (d) Corrected sequence similarity scores are used for graph construction and orthogroup inference. (e) OrthoHMM outputs several files to facilitate downstream analyses. For example, summary files like the gene count per taxon for each orthogroup may be useful in reconstructing gene family evolution, as orthogroups often serve as a proxy for gene families. Additionally, directories of all orthogroups and single-copy orthogroups in FASTA format are created.

To evaluate the efficacy of this approach, we benchmarked OrthoHMM alongside other orthogroup inference methods—OrthoFinder^4,5^, Proteinortho^7^, OrthoMCL^6^, and InParanoid^13,14^, which use BLAST^8^, DIAMOND^9^, or MMseqs^10^ for their sequence similarity search step (see Table S1 for key features of each software). We evaluated OrthoHMM performance using six popular amino acid substitution matrices (Table S2), although more are supported. Our benchmarking used the OrthoBench^15^ dataset, a standardized set of 70 expert-curated reference orthogroups from 12 Bilateria that presents numerous challenges for algorithmic inference (e.g., large multi-copy orthogroups or small rare orthogroups).

During benchmarking, we used strict definitions of true positives, false positives, and false negatives^3,15^. Specifically, true positives were defined as having all the correct members of the orthogroup present and no erroneously included sequences. An orthogroup was determined to be a false positive or false negative if a sequence was erroneously included or excluded, respectively. A predicted orthogroup could be both false positive and false negative if sequences were both erroneously included and orthologous sequences were erroneously excluded (Fig. S1). Similar to other benchmarking studies^4^, true negatives could not be assigned due to only a subset of sequences being assigned to gene families.

This approach is distinct from others, which use ortholog pairs to quantify true positives, false positives, and false negatives. We advocate for whole orthogroup-based benchmarking for at least two reasons: using ortholog pairs inherently overweights large orthogroups and, perhaps more importantly, orthogroups, not gene pairs, are the substrate of downstream analyses (e.g., phylogenomics and inferring gene family gain and loss events). After quantifying these metrics, we calculated three summary performance metrics: precision, recall, and the Fowlkes-Mallows Index (FMI) score, which quantifies the similarity between inferred and reference orthogroups (see supplementary text for details).

Examination of performance metrics revealed that OrthoHMM using specific substitution matrices outperformed all other algorithms (Fig 2a and Table S3). Specifically, OrthoHMM using the BLOSUM90, BLOSUM62, PAM70, or PAM120 substitution matrix for HMM parameterization surpasses the pareto frontier (or the limit of performance) set by previous tools. Next, we integrated the three-performance metrics into a single quantitative measure using rank-based desirability functions. These mathematical operations have been used to combine multiple measures of performance to, for example, benchmark software, rank the potential of chemical compounds for drug discovery, and other applications^16–18^ (see supplementary text for details). In this analysis, the possible scores range from 0 to 1, wherein higher values indicate better algorithm performance. This analysis reinforces OrthoHMM as a top-performing algorithm (Fig. 2b).

**Figure 2.**
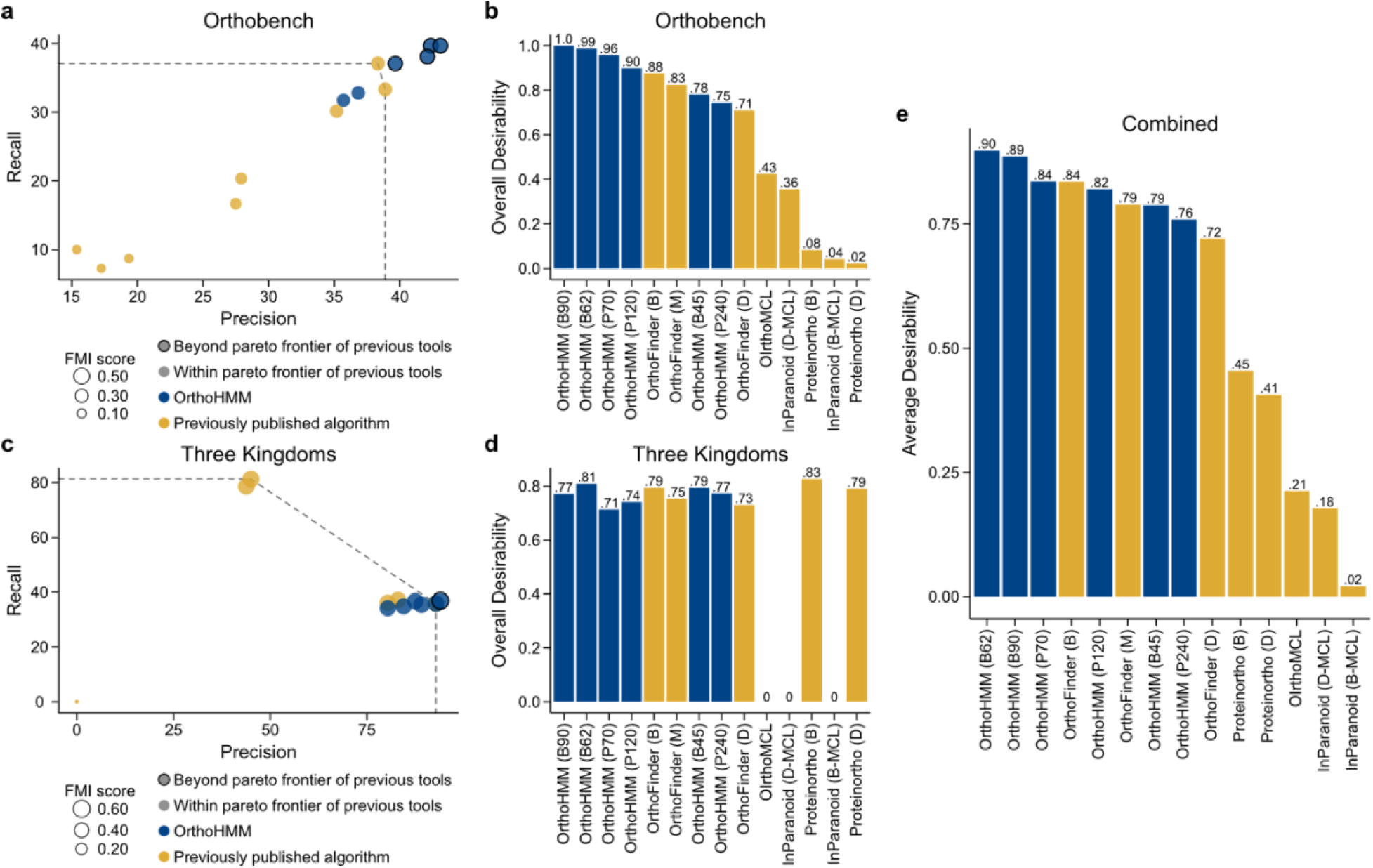
OrthoHMM outperforms other algorithms across two benchmarking datasets. (a) Examination of various algorithms across the three-performance metrics—precision (x-axis), recall (y-axis), and Fowlkes-Mallows Index (FMI) score (data point size)—using the OrthoBench dataset revealed four variants of OrthoHMM outperformed other algorithms. (b) Desirability-based ranking, which aggregates precision, recall, and FMI score, revealed OrthoHMM using the BLOSUM90, BLOSUM62, PAM70, and PAM120 substitution matrices outperformed all other approaches. (c) Examination of the same performance metrics in the Three Kingdoms dataset revealed that one variant of OrthoHMM surpassed the pareto frontier of previous tools. (d) Desirability-based ranking corroborated OrthoHMM is a top-performing algorithm. (e) The average desirability scores in the OrthoBench and Three Kingdoms datasets revealed OrthoHMM with the BLOSUM62 and BLOSUM90 substitution matrices outperformed all other approaches. In panels a and c, the pareto frontier of previous tools is represented as a dashed line. In panels b, d, and e, the abbreviations in the paratheses provide additional information about orthology inference parameters and are as follows: B45, BLOSUM45; B62, BLOSUM62; B90, BLOSUM90; P70, PAM70; P120, PAM120; P240, PAM240; B, BLAST; D, DIAMOND; D-MCL, DIAMOND and Markov clustering; and B-MCL, BLAST and Markov clustering. OrthoHMM is depicted in blue; other algorithms are depicted in gold.

While the OrthoBench dataset is valuable for benchmarking orthology inference methods, benchmarking against another dataset may provide additional insight into the performance of various algorithms. Thus, we engineered a novel dataset for benchmarking orthology inference algorithms, termed the Three Kingdoms dataset. This dataset comprises 255 reference orthologs of near-universally single-copy genes (or BUSCO genes) from 12 proteomes spanning three major eukaryotic kingdoms—four each from animals, plants, and fungi (Table S4). To do so, we obtained BUSCO genes from the eukaryotic database of orthologs^19,20^ and individually identified BUSCO genes in each proteome. The resulting presence, absence, and copy number of BUSCO genes across the 12 proteomes was used to construct reference orthogroups.

While this dataset does have some caveats—for example, only (near) single-copy orthogroups are evaluated, not large multi-copy orthogroups or small rare orthogroups—we suggest that this benchmarking is valuable because single-copy orthogroups are frequently used for downstream analyses like phylogenomics.

Moreover, compared to OrthoBench, this dataset spans a greater phylogenetic distance, an increasingly common feature of many comparative studies, and it is more challenging to infer orthology across deep timescales due to higher levels of sequence divergence between orthologs^21–23^.

Examination of the performance metrics in the Three Kingdoms dataset revealed OrthoHMM using the BLOSUM62 substitution matrix exceeded the pareto frontier set by previous algorithms (Fig. 2c). Desirability-based ranking revealed that most algorithms performed similarly, except for OrthoMCL and InParanoid, which were likely challenged by the phylogenetic distance represented in the Three Kingdoms dataset (Fig. 2d). The top two performing algorithms were Proteinortho using BLAST for sequence similarity search and OrthoHMM using the BLOSUM62 substitution matrix for HMM parameterization (desirability scores of 0.83 and 0.81, respectively). Excluding OrthoMCL and InParanoid, all other algorithms had desirability scores between 0.71 and 0.79.

Interestingly, some algorithms performed well in one dataset but not the other. For example, Proteinortho using either BLAST or DIAMOND for sequence similarity was a top-performing algorithm in the Three Kingdoms dataset, but not the Orthobench dataset. This observation may suggest that Proteinortho can identify near-single copy orthogroups robustly but is challenged by more complex orthogroups (e.g., large multi-copy orthogroups) represented in the Orthobench dataset. In contrast, methods like OrthoHMM and OrthoFinder performed more consistently in both datasets.

Ideally, an algorithm will perform well across diverse evolutionary scenarios and experimental designs. Thus, to rank algorithm performance across both datasets, we took the average overall desirability scores from the OrthoBench and Three Kingdoms datasets for each algorithm. This analysis revealed that four of the top five performing approaches were OrthoHMM-based, indicating that OrthoHMM frequently outperformed other approaches. Notably, two OrthoHMM-based approaches, which used the OrthoHMM using the BLOSUM62 and BLOSUM90 substitution matrices for HMM parameterization, outperformed other algorithms (Fig. 2e). The next best-performing algorithms were OrthoHMM using the PAM70 substitution matrix for HMM parameterization and OrthoFinder using BLAST for sequence similarity search, followed by OrthoHMM using the PAM120 substitution matrix for HMM parameterization.

We next examined the impact of algorithm choice on common downstream analyses. For example, inferring gene family gain and loss patterns in a phylogeny of the genus *Saccharomyces* (Fig. S2a) revealed extensive, software-specific variation (Fig. S2b-c). Similarly, software-specific variation was observed when defining the pangenome of *Candida auris*, a microbial pathogen (Fig. S2d). These analyses suggest that algorithm choice affects the results of downstream analyses and underscore the importance of choosing accurate algorithms.

In summary, benchmarking with empirical data demonstrates that OrthoHMM is a top-performing algorithm for inferring accurate orthogroups. Our benchmarking results indicate that OrthoHMM with the BLOSUM90 or BLOSUM62 data matrices perform the best. We suggest using the BLOSUM90 (or BLOSUM62) matrix if the anticipated divergence between two proteomes is approximately 90% (or 62%). We anticipate OrthoHMM will help facilitate accurate comparative and evolutionary studies, such as refining our understanding of how genes and genomes evolve and the Tree of Life. OrthoHMM is available as an open-source software available for download on PyPi (https://pypi.org/project/orthohmm) and GitHub (https://github.com/JLSteenwyk/orthohmm), complete with documentation (https://jlsteenwyk.com/orthohmm). OrthoHMM dependencies include NumPy^24^, HMMER^11^ (http://hmmer.org/), and MCL^25^ (https://github.com/micans/mcl).

## Supplementary figures, legends, and tables

**Fig. S1.**
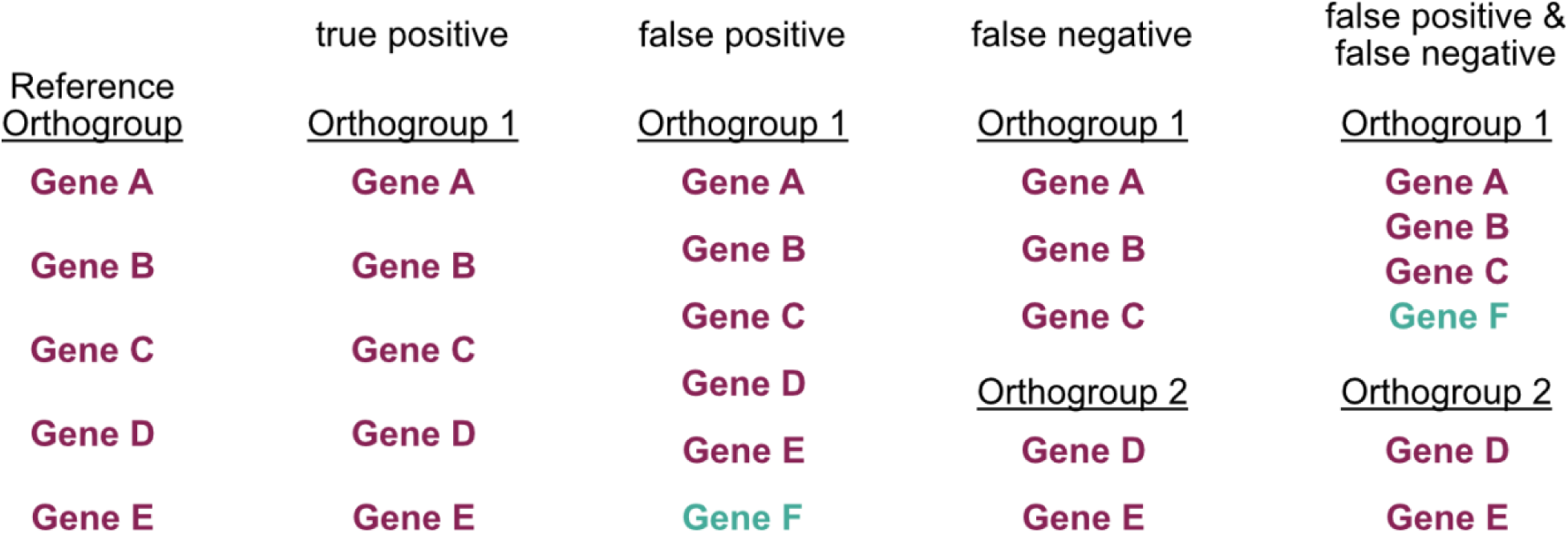
Defining true positives, false positives, and false negatives. Each reference orthogroup is compared to orthogroups inferred by the benchmarked algorithms. An orthogroup will be counted as a true positive if all genes in the reference orthogroup (maroon) are exclusively present in the inferred orthogroup. An orthogroup will be counted as a false positive if an additional gene has been erroneously included (teal). A false negative is when a reference orthogroup is erroneously split across two or more orthogroups. Lastly, a reference orthogroup can be counted as both a false negative and false positive if the algorithm erroneously included genes and split the reference genes across multiple orthogroups.

**Fig. S2.**
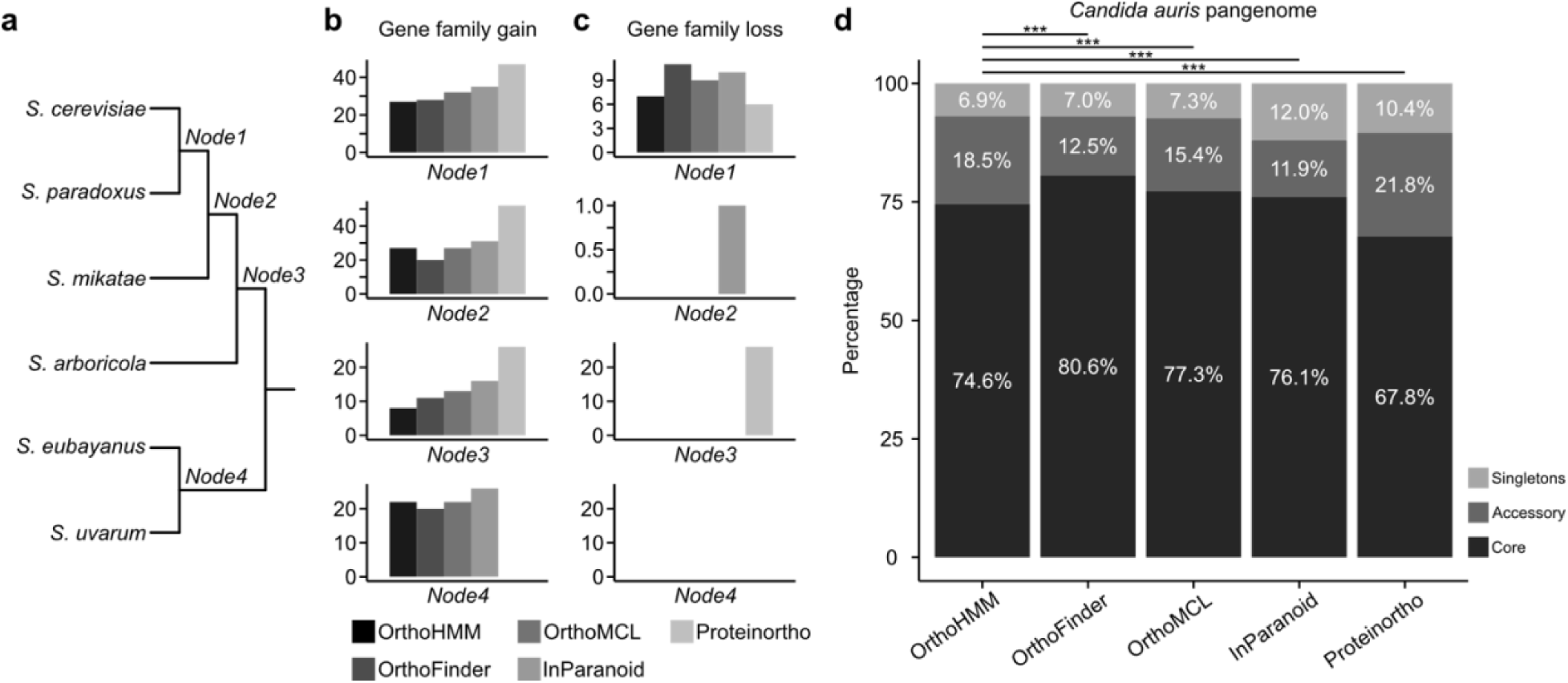
The influence of algorithm choice on common genome-scale analyses that use orthogroup inferences. (a) To evaluate the impact of algorithm choice on gene family evolution, gene family gain and loss were evaluated along ancestral nodes from species in the *Saccharomyces* genus. Different algorithms varied in the number of gene families that were (b) gained or (c) lost. For example, at Node 3—the ancestor of *S. cerevisiae*, *S. paradoxus*, *S. mikatae*, and *S. arboricola*—Proteinortho inferred the most gains whereas OrthoHMM inferred the fewest. (d) Similarly, algorithms varied in the percentage of gene families in the pangenome of a fungal pathogen, *Candida auris*, that was core (present in all isolates; dark grey), accessory (present in two or more isolates, but not all isolates; grey), or singletons (present in a single isolate; light grey). Moreover, the portions significantly varied between OrthoHMM and the other algorithms (p<0.001 for all comparisons, Fisher’s exact test with Benjamini-Hochberg multi-test correction). For each software, only the best-performing iteration was used. At these evolutionary distances, OrthoHMM using the BLOSUM90 substitution matrix was used since it performed the best in the Bilateria dataset and the evolutionary divergence of the sequences compared is relatively recent^26^. For OrthoMCL, Proteinortho, InParanoid, and OrthoFinder, BLAST was used during sequence similarity search.

**Table S1.** Other algorithms used to evaluate OrthoHMM performance. OrthoHMM was benchmarked across four other software tools. Where possible, multiple different sequence similarity search approaches were tested. For example, when using OrthoFinder, sequence similarity search using BLAST, DIAMOND, and MMseqs was tested.

| Software name and version | Sequence similarity search method | Corrects for sequence length bias | Corrects for phylogenetic distance | Clustering method | Reference |
| --- | --- | --- | --- | --- | --- |
| Proteinortho, v6.3.2 | BLAST | No | No | Spectral partitioning | 7 |
|  | DIAMOND |  |  |  |  |
| InParanoid, v5.0 | BLAST | No | No | Markov clustering | 13,14 |
|  | DIAMOND |  |  |  |  |
| OrthoMCL, v1.4 | BLAST | No | No | Markov clustering | 6 |
| OrthoFinder, v2.2.7 | BLAST | Yes | Yes | Markov clustering | 4,5 |
|  | DIAMOND |  |  |  |  |
|  | MMseqs |  |  |  |  |
| OrthoHMM | HMMER | Yes | Yes | Markov clustering | This study |

**Table S2.** Amino acid substitution matrices tested for HMM parameterization in OrthoHMM. The matrices supported in OrthoHMM include BLOSUM and PAM matrices. Each matrix is followed by a number (e.g., 62 in BLOSUM62). In the case of the BLOSUM matrix, lower values indicate greater evolutionary distances captured by the matrix. In the case of the PAM matrix, lower values indicate relatively shorter evolutionary. Thus, PAM240 is more analogous to BLOSUM45 than it is to BLOSUM90. PAM is an abbreviation of Point Accepted Mutation. BLOSUM refers to BLOcks SUbstitution Matrix.

| Substitution matrix | Description | Used in Benchmarking |
| --- | --- | --- |
| BLOSUM90 | Built from sequences clustered at 90% similarity | Yes |
| BLOSUM62 | Built from sequences clustered at 62% similarity | Yes |
| BLOSUM45 | Built from sequences clustered at 45% similarity | Yes |
| PAM240 | Built from sequences that have 240 mutations per 100 amino acids | Yes |
| PAM120 | Built from sequences that have 120 mutations per 100 amino acids | Yes |
| PAM70 | Built from sequences that have 70 mutations per 100 amino acids | Yes |
| BLOSUM80 | Built from sequences clustered at 80% similarity | No |
| BLOSUM50 | Built from sequences clustered at 50% similarity | No |
| PAM30 | Built from sequences that have 30 mutations per 100 amino acids | No |

**Table S3.** Results from benchmarking using OrthoBench. The results from benchmarking various algorithms are summarized here. This includes the number of correctly inferred orthogroups (true positives), the number false positives, and the number of false negatives. These values were used to calculate recall, precision, F1, and FMI scores. Notably, OrthoHMM with the BLOSUM62 and BLOSUM90 substitution matrices outperformed other approaches. Moreover, OrthoHMM was the only algorithm to have performance metric values that surpassed 0.40. The abbreviations in the paratheses provide additional information about orthology inference parameters and are as follows: B45, BLOSUM45; B62, BLOSUM62; B90, BLOSUM90; P70, PAM70; P120, PAM120; P240, PAM240; B, BLAST; D, DIAMOND; D-MCL, DIAMOND and Markov clustering; and B-MCL, BLAST and Markov clustering.

|  | Proteinortho<br>(B) | Proteinortho<br>(D) | InParanoid<br>(B-MCL) | InParanoid<br>(D-MCL) | OrthoMCL | OrthoFinder<br>(B) | OrthoFinder<br>(D) | OrthoFinder<br>(M) | OrthoHMM<br>(P70) | OrthoHMM<br>(P120) | OrthoHMM<br>(P240) | OrthoHMM<br>(B45) | OrthoHMM<br>(B62) | OrthoHMM<br>(B90) |
| --- | --- | --- | --- | --- | --- | --- | --- | --- | --- | --- | --- | --- | --- | --- |
| True (+) | 6 | 5 | 9 | 11 | 12 | 23 | 19 | 21 | 24 | 23 | 20 | 21 | 25 | 25 |
| False (+) | 25 | 23 | 33 | 29 | 31 | 37 | 35 | 33 | 33 | 35 | 36 | 36 | 34 | 33 |
| False (-) | 64 | 65 | 54 | 55 | 47 | 39 | 44 | 42 | 39 | 39 | 43 | 43 | 38 | 38 |
| Recall | 0.09 | 0.07 | 0.14 | 0.17 | 0.2 | 0.37 | 0.3 | 0.33 | 0.38 | 0.37 | 0.32 | 0.33 | 0.4 | 0.4 |
| Precision | 0.19 | 0.18 | 0.21 | 0.28 | 0.28 | 0.38 | 0.35 | 0.39 | 0.42 | 0.4 | 0.36 | 0.37 | 0.42 | 0.43 |
| F1 score | 0.12 | 0.1 | 0.17 | 0.21 | 0.23 | 0.37 | 0.32 | 0.36 | 0.4 | 0.38 | 0.34 | 0.35 | 0.41 | 0.41 |
| FMI score | 0.13 | 0.11 | 0.17 | 0.22 | 0.24 | 0.37 | 0.32 | 0.36 | 0.4 | 0.38 | 0.34 | 0.35 | 0.41 | 0.41 |

**Table S4.** Eukaryotic organisms in the Three Kingdoms dataset. The Three Kingdoms dataset comprises four proteomes from the three eukaryotic kingdoms— animals, plants, and fungi. These species were selected for their status as model (or emerging model) organisms that span much of the diversity of each kingdom.

| Kingdom | Common name | Genus and species | NCBI Accession |
| --- | --- | --- | --- |
| Animal | Sponge | *Amphimedon queenslandica* | GCF_000090795.2 |
| Animal | Green anole | *Anolis carolinensis* | GCF_035594765.1 |
| Animal | Zebrafish | *Danio rerio* | GCF_000002035.6 |
| Animal | African clawed frog | *Xenopus laevis* | GCF_017654675.1 |
| Plant | Thale cress | *Arabidopsis thaliana* | GCF_000001735.4 |
| Plant | Rice | *Oryza sativa* | GCF_034140825.1 |
| Plant | Spikemoss | *Selaginella moellendorffii* | GCF_000143415.4 |
| Plant | Corn | *Zea mays* | GCF_902167145.1 |
| Fungi | Filamentous fungus | *Aspergillus fumigatus* | GCF_000002655.1 |
| Fungi | Baker's or brewer's yeast | *Saccharomyces cerevisiae* | GCF_000146045.2 |
| Fungi | Splitgill mushroom | *Schizophyllum commune* | GCF_000143185.2 |
| Fungi | Fission yeast | *Schizosaccharomyces pombe* | GCF_000002945.2 |

**Table S5.** Results from benchmarking using the Three Kingdoms Dataset. The results from benchmarking various algorithms are summarized here. This includes the number of correctly inferred orthogroups (true positives), the number false positives, and the number of false negatives. These values were used to calculate recall, precision, F1, and FMI scores. The abbreviations in the paratheses provide additional information about orthology inference parameters and are as follows: B45, BLOSUM45; B62, BLOSUM62; B90, BLOSUM90; P70, PAM70; P120, PAM120; P240, PAM240; B, BLAST; D, DIAMOND; D-MCL, DIAMOND and Markov clustering; and B-MCL, BLAST and Markov clustering.

|  | Proteinortho<br>(BLAST) | Proteinortho<br>(DIAMOND) | InParanoid<br>(BLAST+MCL) | InParanoid<br>(DIAMOND+MCL) | OrthoMCL | OrthoFinder<br>(BLAST) | OrthoFinder<br>(DIAMOND) | OrthoFinder<br>(MMSeqs) | OrthoHMM<br>(PAM70) | OrthoHMM<br>(PAM120) | OrthoHMM<br>(PAM240) | OrthoHMM<br>(BLOSUM45) | OrthoHMM<br>(BLOSUM62) | OrthoHMM<br>(BLOSUM90) |
| --- | --- | --- | --- | --- | --- | --- | --- | --- | --- | --- | --- | --- | --- | --- |
| True (+) | 113 | 110 | 0 | 0 | 0 | 91 | 90 | 93 | 86 | 87 | 91 | 91 | 94 | 90 |
| False (+);<br>overclumped | 26 | 30 | 123 | 122 | 18 | 163 | 159 | 157 | 166 | 163 | 157 | 163 | 161 | 164 |
| False (-);<br>oversplit | 138 | 141 | 146 | 142 | 254 | 7 | 22 | 19 | 21 | 16 | 13 | 7 | 6 | 11 |
| Recall | 81.29 | 78.57 | 0 | 0 | 0 | 35.83 | 36.14 | 37.2 | 34.13 | 34.8 | 36.69 | 35.83 | 36.86 | 35.43 |
| Precision | 45.02 | 43.82 | 0 | 0 | 0 | 92.86 | 80.36 | 83.04 | 80.37 | 84.47 | 87.5 | 92.86 | 94 | 89.11 |
| F1 score | 57.95 | 56.26 | 0 | 0 | 0 | 51.71 | 49.86 | 51.38 | 47.91 | 49.29 | 51.7 | 51.71 | 52.95 | 50.7 |
| FMI score | 60.5 | 58.68 | 0 | 0 | 0 | 57.68 | 53.89 | 55.58 | 52.37 | 54.22 | 56.66 | 57.68 | 58.86 | 56.19 |

## Supplementary text

**Detailed explanation of the OrthoHMM algorithm**

### User arguments

OrthoHMM can be flexibly run and provides numerous user arguments to enable researchers to tailor different OrthoHMM runs to their specifications. The only required argument is the path to the directory of FASTA files that will be used for calling orthogroups. Inside that directory, OrthoHMM will automatically detect files that end in the suffix *.fa*, *.faa*, *.fas*, *.fasta*, *.pep*, or *.prot*. A full list of OrthoHMM user arguments are as follows:

1. -o, --output_directory: the path to the output directory to store OrthoHMM results (default: same directory as the input files);
2. -p, --phmmer: the path to the phmmer executable from the HMMER suite^11^ (default: assumed to be part of the user’s PATH variable; i.e., phmmer can be evoked by typing ‘phmmer’);
3. -e, --evalue: expectation value (or e-value) threshold used when filtering phmmer results (default: 0.001);
4. -x, --substitution_matrix: substitution matrix used for creating the profile HMM (default: BLOSUM62 and additionally supported substitution matrices include BLOSUM45, BLOSUM50, BLOSUM80, BLOSUM90, PAM30, PAM70, PAM120, and PAM240);
5. -c, --cpu: the number of CPU workers for multithreading during sequence search and other file processing steps (default: the number of CPUs available will be auto-detected);
6. -s, --single_copy_threshold: taxon occupancy threshold for single-copy orthologous genes specified as a fraction (default: 0.5 or 50% occupancy threshold);
7. -m, --mcl: path to mcl executable^25^ (default: assumed to be part of the user’s PATH variable; i.e., mcl can be evoked by typing ‘mcl’);
8. -i, --inflation_value: Markov clustering inflation parameter for clustering genes into orthogroups (default: 1.5; lower values are more permissive resulting in larger orthogroups, whereas higher values are stricter resulting in smaller orthogroups);
9. --stop: internal OrthoHMM step to stop at (default: write; other choices include prepare [stop after preparing input files for the all-by-all search]; infer [stop after inferring orthogroups]; and write [stop after writing sequence files for the orthogroups]); and
10. 10.--start: internal OrthoHMM step to start analysis at (default: none, which will start OrthoHMM from the beginning of the pipeline; other options include search_res, which will start the analysis after the all-by-all search has completed).

The stop-and-start steps implemented in OrthoHMM can help users pseudo-parallelize parts of the OrthoHMM workflow. For example, it may be faster to spread the all-by-all search across diverse nodes in a high-performance computing cluster. To do so, users can have OrthoHMM print the all-by-all search commands by setting the “stop” argument to “prepare.” Next, users can run the all-by-all searchers independent of OrthoHMM. Once completed, users can run OrthoHMM with the “start” parameter set to “search_res” to start the OrthoHMM workflow from completed all-by-all search files.

### OrthoHMM workflow

Five steps summarize key features of OrthoHMM’s internal workflow. The first step is executing the all-by-all search using phmmer from the HMMER suite^11^. At the time of development, OrthoHMM was tested using the latest stable HMMER release, version 3.4. During sequence similarity search, every ordered pairwise combination is conducted

In the second step, the resulting sequence similarity scores are used to determine network edges between every pairwise combination of taxa. This requires defining thresholds for significant edges. To do so, sequence search results are first filtered by an e-value threshold. Next, scores are corrected by gene lengths by taking the score and dividing it by the sum of the query and target gene lengths. The resulting scores are then phylogenetically corrected using the average reciprocal best hit score between the two taxa. The resulting information sets gene-specific thresholds for every pairwise combination of taxa.

In the third step, these thresholds are used to determine significant network edges. Network edges with corrected values greater than or equal to the threshold are kept, while putative edges with corrected values lower than the threshold are discarded. All retained edges are then outputted to an edges file.

In the fourth step, Markov clustering is conducted using the MCL program^25^. Following established protocol and benchmarking^4,6,27^, an inflation parameter of 1.5 is used. The inflation parameter influences the “tightness” of resulting clusters; lower values will result in more permissive, larger orthogroups, and higher values will result in stricter, smaller orthogroups. During OrthoHMM development, MCL version 22-282 was used.

In the fifth step, OrthoHMM generates all output files. This includes the summary files and FASTA files for each orthogroup. These files aim to facilitate downstream analysis, such as charting gene family gain and loss and phylogenomic inference.

### Ensuring long-term stability

Approximately 30% of all bioinformatic tools fail to install^28^, reflecting a lack of long-term software stability. To ensure OrthoHMM has long-term stability, we followed software design and development practices that followed industry standards and other software we engineered (e.g., ClipKIT and PhyKIT^18,29^). Specifically, the OrthoHMM codebase features a modular design to facilitate easy debugging and integrating new features. We also wrote numerous unit and integration tests that test the efficacy of the codebase.

Lastly, we implemented a continuous integration pipeline that automatically tests OrthoHMM across different Python versions and ensures expected outputs are produced. Together, these features will help ensure that OrthoHMM is a longstanding bioinformatic tool for the research community.

#### Other ortholog calling algorithms and parameters used

Eight ortholog calling algorithms (excluding OrthoHMM) were employed to determine if OrthoHMM improves over other approaches. Other algorithms include Proteinortho^7^, InParanoid^13,14^, OrthoMCL^6^, and OrthoFinder^4,5^ (Table S1). For Proteinortho and InParanoid, BLAST or DIAMOND were used for the sequence similarity search. For OrthoFinder, BLAST, DIAMOND, or MMseqs was used for sequence similarity search. For OrthoMCL, BLAST was used for sequence similarity search. Because InParanoid does not infer orthogroups, but instead infers ortholog pairs, subsequent Markov clustering^25^ was performed from InParanoid output using an inflation value of 1.5.

Excluding which algorithm was used for sequence similarity search, all other arguments were set to default values. Combined with the six variations of OrthoHMM that were used for benchmarking, 14 total ortholog calling algorithms were employed.

#### Calculating performance metrics

To evaluate the performance of each algorithm, whole orthogroup definitions of true positive, false positive, and false negatives were used (Fig. S1). After determining the number of true positives and false positives and negatives, precision, recall, FMI, and F1 scores were calculated. Precision represents the proportion of the positive classifications that are correct and is calculated using the following formula:

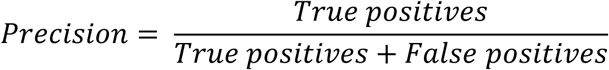

Recall is the proportion of all positives that were correct and is calculated using the following formula:

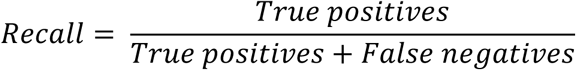

To evaluate the balance between precision and recall, the FMI and F1 scores were also calculated. The FMI score is the square root of the product of precision and recall and is calculated using the following formula:

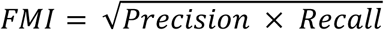

The F1 score is the harmonic mean of precision and recall and is calculated using the following formula:

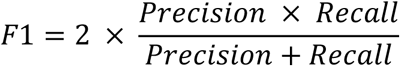

While we calculated both the FMI and F1 score, we present only the FMI score in the main text but show the F1 score values in the supplementary information for completeness (see Table S2 and S4).

To integrate these values into a single rank-based metric, we used desirability functions. Desirability functions facilitate incorporating diverse data types into a single metric and have been used to rank compounds for their potential as clinical drugs, the association of genes to various phenotypes like preterm birth, and the performance of different algorithms^16–18^. To do so, precision, recall, and FMI scores were individually scaled to be between 1 (best performing) and 0 (worst performing) using the following formula:

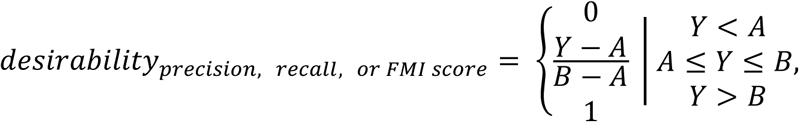

where *Y* is the variable value (precision, recall, or FMI score), *A* is the minimum variable value across all algorithms, and *B* is the maximum variable value across all algorithms. In this way, the best performing algorithm got a desirability score of 1 and the worst performing algorithm got a desirability score of 0. After individually scaling precision, recall, or FMI scores to desirability scores, the overall desirability for each dataset was calculated by taking the average desirability score for each metric. To integrate desirability scores from the OrthoBench and Three Kingdoms dataset, the average score was taken from the overall desirability scores.

#### Exemplary comparative and evolutionary genomic analysis

To better understand the influence of algorithm selection on common analyses downstream of orthology inference, gene family gain and loss analysis as well as defining the core and accessory pangenomes was conducted. To conduct gene family gain and loss analysis, the respective outputs were converted into a matrix with rows representing orthogroups (a common proxy for gene families) and columns representing the distinct species. The resulting matrix was used to infer ancestral gain and loss events using Dollo parsimony with the software Count, v10.04^30^. This analysis was conducted using publicly available proteomes of *Saccharomyces cerevisiae* (GCF_000146045.2), *S. paradoxus* (GCF_002079055.1), *S. mikatae* (GCF_947241705.1), *S. arboricola* (GCA_000292725.1), *S. eubayanus* (GCF_001298625.1), and *S. uvarum* (GCA_947243795.1) from NCBI. Default parameters were used for each algorithm.

To determine the influence of algorithm choice on how pangenomes are defined, we conducted a similar analysis among all publicly available proteomes of *Candida auris* (NCBI accessions: GCA_002759435.3, GCA_003014415.1, GCA_007168705.1, GCA_008275145.1, GCA_016772135.1, GCA_016809505.1, GCA_019332045.1, GCA_041381755.1, and GCF_003013715.1). Using the output of each algorithm, we determined the proportion of gene families that were core (present in all isolates), accessory (present in fewer than 100% of isolates but more than one), or singletons (present in only one isolate). To determine if the proportions of each category varied between orthology inference approaches, a Fisher’s exact test was used with Benjamini-Hochberg multi-test correction.

## Acknowledgements

JLS is a Howard Hughes Medical Institute Awardee of the Life Sciences Research Foundation. Research in the King lab is supported by the Howard Hughes Medical Institute. We thank the King lab for the helpful discussion and comments. We also thank personnel from the Hittinger and Rokas labs for helpful feedback. We thank Dr. Sean Eddy for providing additional feedback.

